# Fitness of reciprocal F_1_ hybrids between *Rhinanthus minor* and *Rhinanthus major* under controlled conditions and in the field

**DOI:** 10.1101/500454

**Authors:** Renate A. Wesselingh, Šárka Hořčicová, Khaled Mirzaei

**Affiliations:** Biodiversity Research Centre, Earth and Life Institute, UCLouvain, Croix du Sud 4 box L7.07.04, B-1348 Louvain-la-Neuve, Belgium; Department of Botany, Faculty of Science, University of South Bohemia, CZ-370 05, České Budějovice, Czech Republic

**Keywords:** emergence, germination, greenhouse, field transplant, hybridization, seed production, stratification, survival

## Abstract

The performance of first-generation hybrids determines to a large extent the long-term outcome of hybridization in natural populations. F_1_ hybrids can facilitate further gene flow between the two parental species, especially in animal-pollinated flowering plants. We studied the performance of reciprocal F_1_ hybrids between *Rhinanthus minor* and *R. major*, two hemiparasitic, annual, self-compatible plant species, from seed germination to seed production under controlled conditions and in the field. We sowed seeds with known ancestry outdoors before winter and followed the complete life cycle until plant death in July the following season. While germination under laboratory conditions was much lower for the F_1_ hybrid formed on *R. major* compared to the reciprocal hybrid formed on *R. minor*, this difference disappeared under field conditions, pointing at an artefact caused by the experimental conditions during germination in the lab rather than at an intrinsic genetic incompatibility. Both F_1_ hybrids performed as well as or sometimes better than *R. minor*, which had a higher fitness than *R. major* in one of the two years in the greenhouse and in the field transplant experiment. The results confirm findings from naturally mixed populations, where F_1_ hybrids appear as soon as the two species meet and which leads to extensive advanced-hybrid formation and introgression in subsequent generations.

## Introduction

Speciation without a change in chromosome numbers, homoploid speciation, is a slow process that initially leaves the door wide open for mating and offspring production with sister species. This is why isolation, i.e. allopatry or some other form of prezygotic isolation, such as divergence in mating preferences, host species, phenology or pollinator guild, is generally considered necessary to complete the speciation process (Abbott *et al.*, 2013). When nascent sister species meet in sympatry and prezygotic isolation is not complete, natural hybridization can occur. The first step in hybridization is the formation of hybrid offspring by interspecific sperm or pollen transfer and subsequent fertilization. If this leads to the production of at least partly viable first-generation or F_1_ hybrids, the outcome of hybridization will strongly depend on the fitness of these hybrids. A strongly reduced fitness for F_1_ hybrids precludes any advanced-hybrid formation or introgression and can be a severe bottleneck. But F_1_ hybrids, both intra-and interspecific, are also known to exhibit heterosis (Birchler *et al.*, 2010) or transgressive trait values, beyond the expected mid-parent value (Rieseberg *et al.*, 1999; Johnston *et al.*, 2004). Once established, even just a few F_1_ hybrids can serve as a bridge to the formation of advanced hybrids and introgression (Arnold *et al.*, 2012), often facilitating pollen transport between the parental species in animal-pollinated angiosperms (Leebens-Mack & Milligan, 1998; Emms & Arnold, 2000). Knowledge about F_1_ fitness is thus crucial for understanding the composition of mixed populations and to predict their future.

Hybrid formation and fitness in flowering plants can vary depending on which species is the maternal parent and lead to asymmetries in fitness between the reciprocal crosses, likely to be caused by Dobzhansky-Muller incompatibilities (Tiffin *et al.*, 2001). The asymmetry in postmating reproductive isolation has been called Darwin’s corollary to Haldane’s rule (Turelli & Moyle, 2007), and interactions between nucleus and cytoplasm, between gametophyte (pollen) and sporophyte (stigma and style), and within the triploid endosperm are common causes of asymmetries in reproductive isolation in angiosperms (Turelli & Moyle, 2007).

While asymmetries in hybrid fitness are often caused by intrinsic factors, hybrid fitness can also be dependent on the environment. Hybrids can have a relatively low fitness in the parental habitats, but perform better in alternative, unique habitats (Cruzan & Arnold, 1993; Gramlich *et al.*, 2016). This can lead to homoploid hybrid speciation if hybrids are (spatially) isolated from their parental lines (Arnold, 1993; Abbott *et al.*, 2013). By growing plants under optimal (greenhouse or growth room) conditions, excluding environmental factors, intrinsic genetic incompatibilities between the parental genomes leading to poor hybrid performance can be identified, as well as possible asymmetries in performance between reciprocal F_1_ hyrids. Field transplants of hybrids of known descent and the parental species allow for the quantification of hybrid fitness (Arnold & Hodges, 1995; Martin *et al.*, 2006; Gramlich & Hörandl, 2016; Favre *et al.*, 2017). Using genetic tools, the prevalence of hybrids in different life stages (seed, seedling, juvenile, adult) can be determined in naturally mixed populations to see if it decreases as a result of consistent selection against hybrids (Cornman *et al.*, 2004; Lindtke *et al.*, 2014; Hipperson *et al.*, 2016). Experiments in which environmental factors are independently varied in a controlled fashion can then help to identify the factors directly responsible for fitness differences (Johnston *et al.*, 2001; Favre & Karrenberg, 2011; Hipperson *et al.*, 2016).

Although knowledge on the fitness of hybrids is crucial for understanding the possible outcomes of hybridization, relatively few studies (Emms & Arnold, 1997; Burke *et al.*, 1998; Kimball *et al.*, 2008; Favre & Karrenberg, 2011; Lepais *et al.*, 2013; Favre *et al.*, 2017) have compared the performance of reciprocal F_1_ hybrids under both controlled conditions and in the field. Some of these studies used seedlings, rooted cuttings or rhizomes for field transplants. This gives the advantage of being able to replicate the same genotype in several environments, but the downside is that part of the life cycle, from seed to established young plant, is missing. This can be justified for long-lived species, but for annual species, including the seed stage is crucial in understanding local adaptation (Postma & Ågren, 2018).

Our study system comprises two annual hemiparasitic plant species, *Rhinanthus minor* L. and *Rhinanthus major* Ehrh. (Orobanchaceae). They are both pollinated by bumblebees (Kwak, 1978; Natalis & Wesselingh, 2012a) and are known to readily hybridize in nature, the hybrid has been described as *Rhinanthus* × *fallax* (Wimm. & Grab.) Chabert (Kwak, 1980). We have studied the composition of mixed populations (Ducarme *et al.*, 2010), possible prezygotic barriers such as bumblebee preference and constancy (Natalis & Wesselingh, 2012a, 2013) and pollen tube growth rates (Natalis & Wesselingh, 2012b). Hybrid seed production after hand pollination with heterospecific pollen is known to be lower on *R. major* than on *R. minor* (Kwak, 1979; Campion-Bourget, 1980a; b; Ducarme & Wesselingh, 2013), and both species produce less hybrid seeds than expected after pollination with a 50:50 pollen mix (Natalis & Wesselingh, 2012b). Despite the wealth of knowledge on hybridisation in this species pair, no study has ever attempted to quantify hybrid fitness in the field, apart from one study (Kwak, 1980) that looked at the number of seeds per flower only, without considering plant size and total seed production, the latter being the fitness measure that really counts.

*Rhinanthus* seeds can only germinate after several weeks of cold stratification (Westbury, 2004; ter Borg, 2005), which limits germination to early spring. Under laboratory conditions, with seeds placed on moist filter paper in petri dishes in a refrigerator at ± 5°C, a strong difference in germination rate is repeatedly observed between the reciprocal F_1_ hybrids. Hybrids formed on *R. major* (F_1a_ hybrids) germinate at rates between 5 and 30%, while F_1m_ hybrids, which have *R. minor* as the maternal parent, germinate as well as or better than *R. minor*, with germination percentages close to 100% (Kwak, 1979; Campion-Bourget, 1980a; Natalis & Wesselingh, 2012b; Ducarme & Wesselingh, 2013). However, it has never been tested if this difference in germination rate also occurs under field conditions.

We therefore set out to record the process of germination of hybrid seeds in the laboratory and to compare performance along the complete life cycle (germination/emergence, survival, seed production) of reciprocal F_1_ hybrids and the parental lines under both greenhouse and field conditions.

In contrast to other study systems, in which the parental species have distinct ecological niches (Campbell *et al.*, 1997; Favre & Karrenberg, 2011; Cahenzli *et al.*, 2018) and transplants can be performed in habitats that are clearly attributed as typical for one of the two parental species, our two study species can co-occur in a range of different grassland types, and only subtle differences in nutrient status seem to determine which of the two will become dominant (Ducarme & Wesselingh, 2010). We therefore included a fertilizer addition treatment in the field experiment. It is known that in nutrient-rich grasslands, where plant growth is very vigorous, *Rhinanthus* seedlings have difficulties establishing themselves in the dense sward due to a lack of light at ground level (Těšitel *et al.*, 2011), but the surviving parasites can profit from the increased nutrient availability for their host by producing more biomass, flowers and seeds (Mudrák *et al.*, 2013). We wanted to investigate the role of grassland nutrient status in determining the relative fitness of the parental species and their hybrids.

We aimed at answering the following questions:

1. Are there differences in performance (germination/emergence, survival, seed production) between the reciprocal F_1_ hybrids and between hybrids and the parental lines under the conditions used in the laboratory and in the field?
2. Is the relative performance of the parental and hybrid classes different between the laboratory and the outdoor conditions?
3. Is there an influence of fertilizer addition on the relative performance of the parental and hybrid classes in the field?

## Materials and methods

### Study species

*Rhinanthus minor L.* and *Rhinanthus major* Ehrh. (Orobanchaceae) are hemiparasitic annual plants occurring in grasslands. Like the other species in the genus *Rhinanthus* they are capable of parasitising a wide range of host species in the Poaceae and Fabaceae. *Rhinanthus major* (synonym *R. angustifolius* C.C. Gmel) is distributed throughout temperate and boreal/alpine Europe, ranging from central Scandinavia to Italy and from France to Russia (von Soó & Webb, 1972). The range of *Rhinanthus minor* overlaps with that of *R. major* and extends further out to the west (the British Isles, Iceland, Greenland and North America), to the north of Scandinavia and to the south (Spain, Corsica, Italy, Greece). All *Rhinanthus* species produce seeds in summer, which stay dormant in the soil until after cold stratification (ter Borg, 2005). Germination in temperate regions starts in February-March and seedlings emerge shortly after. Flowering usually starts in May in the vernal ecotype of both species (von Soó & Webb, 1972), which is adapted to hay making by flowering early and at a relatively small size. By early July most seeds have ripened and capsules dehisce. The heavy seeds (2-3 mg: Westbury, 2004) fall out of the capsule when the dead stalks break or are mown, but they can be dispersed over longer distances by mowing machinery (Strykstra *et al.*, 1996) or in the hay itself (Vrancken *et al.*, 2012).

### Hybrid production

The general procedure to produce *Rhinanthus* hybrids in our lab is to collect seeds in pure populations in July, keep them dry and cool in order to prolong seed longevity until October-November and germinate the seeds in petri dishes in a refrigerator (± 5-7°C). The emerging seedlings are then planted in pots with host plants (*Trifolium repens*) in a heated greenhouse in January-February and crosses are made by hand pollination when plants start to flower, which is around two months after planting. The capsules are harvested when dry (± 3 weeks after pollination) and the number of seeds per capsule is counted. The dry seeds are then stored in closed recipients in a refrigerator until sowing in autumn for the next greenhouse generation. The specific details for each experiment are given below.

For the transplant experiment in 2013–2014, seeds were collected in pure populations of each species. The source populations for *R. minor* were the nature reserve Housta-Darquenne (July 2010) and the local population on the campus of UCLouvain (July 2011), which had been sown in 2003 using seeds from this nature reserve. For *R. major*, seeds were collected in the nature reserve Doode Bemde in July 2010 and in a local population on the UCLouvain campus in July 2012, which had been sown in 2003 using seeds collected in the Doode Bemde population.

Fifty seeds per population were put in petri-dishes on moist filter paper on 31 October 2012 and stored in a refrigerator at around 5°C. Germination started after ± 4 weeks for the seeds collected in 2010 and 2011, and after 8 weeks for the seeds collected in 2012. The seedlings were kept in the refrigerator until the cotyledons emerged from the seed coat and planted in pots with a single host plant (*Trifolium repens*), which had been sown on 24 October 2012 in a heated greenhouse in 0.75 L square pots (10 x 10 cm surface area). Each corner of a pot received one seedling, so a pot was occupied by maximum 4 plants, and pots only contained plants from the same population. Flowering started in the greenhouse on 12 March 2013, and crosses were made by hand pollination preceded by emasculation of the closed bud (only needed on *R. minor*) to prevent autonomous self-pollination (Ducarme & Wesselingh, 2013). We produced hybrid seeds in both directions as well as pure seeds by performing intraspecific crosses (including selfing). After the fruits ripened and started to dehisce in March-May 2013, the capsules were harvested and left to dry in 24-well plates. After counting the number of seeds produced per fruit, the closed plates were kept in a refrigerator until the start of the experiments.

### Germination under controlled conditions

The seeds that were produced in the greenhouse in spring 2013 and that were not used in the field transplants (see Performance in the field) were put in small petri dishes on moist filter paper (one dish per cross) and placed in a refrigerator at 5°C on 23 October 2013 for the production of F_1_ and F_2_ and backcross hybrids in the greenhouse. The number of seeds per cross ranged from 1 to 11, with an average of 5.2 seeds per cross, with in total 72 *R. major* seeds, 77 *R. minor* seeds, 106 F_1m_ hybrid seeds and 106 F_1a_ hybrid seeds. From 29 November 2013 onwards, germination was checked at least once a week until 5 March 2014. Seeds with a protruding radicle were considered as germinating and put to one side in each petri dish to facilitate subsequent checks.

### Greenhouse performance

Mortality in the greenhouse is generally very low (we typically lose less than 5% of the seedlings after planting) and pollination is done manually and with different pollen sources, leading to differences among plants in seed production. We therefore scored performance in the greenhouse using flower production, which is a very good proxy of plant biomass (Ducarme & Wesselingh, 2010) and seed production under natural pollination (see Performance in the field) and hence fitness in *Rhinanthus*. We recorded flower production for the parents of the hybrids (14 *R. minor* and 11 *R. major*) in the greenhouse in spring 2013 together with a group of simultaneously grown F_1_ hybrids (24 F_1m_ and 12 F_1a_) that had been produced in the greenhouse in the previous year.

Since all plant growing activities were moved to a new greenhouse in January 2014, we repeated the experiment in spring 2013 with the seedlings from the germination experiment (see Germination under controlled conditions) that were grown in the new greenhouse to produce new hybrids, but otherwise using the same methods. This time we had 64 *R. minor* and 59 *R. major* plants, issued from intraspecific crosses between the parents, plus 88 F_1m_ and 15 F_1a_ hybrids. In both greenhouses, the temperature was regulated around 20°C in the day and 18°C at night by central heating to increase the temperature and opening the windows to decrease it. The new greenhouse also regulated relative humidity (at 60%) and used LED lights in the photosynthetically active spectrum for illumination (16h daylength), while in the old greenhouse, this was done with mercury vapour lamps.

### Performance in the field

In December 2013, 8 experimental plots of 100 × 50 cm each were set up in a grassland on the UCLouvain campus that had been a lawn until 2009, when the area had been fenced and partly sown with seeds of both *Rhinanthus* species at one end for observations on bumblebee behaviour in 2010 (Natalis & Wesselingh, 2013). Although on loamy soil, the vegetation in this grassland is not very productive, due to decades of regular mowing without any fertilizer addition, and at the time of our experiment, *Festuca rubra* L. was the dominant grass species. We used a total of 1152 seeds (144 per plot) that were produced in the greenhouse (see Hybrid production), of which 188 were *R. major*, 368 *R. minor*, 240 F_1a_ hybrids and 356 F_1m_ hybrids. We made a design that distributed pairs of seeds from the same cross randomly over four plots. Six 96-well plates (8 × 12 wells per plate, 1.5 plates per pair of plots) were filled with moistened white sand and two seeds of the same cross were placed in the sand in each well. The plates were then kept in the refrigerator until planting in the field plots one week later, on 10-11 December 2013. In order to plant the seeds, we placed a grid, made of a piece of fencing with a square 13-mm mesh, in each plot and single seeds were sown 5.25 cm apart (4 cells in the grid) in 8 rows and 18 columns by making a 1-cm deep hole in the middle of the grid cell with a wooden stick and dropping the seed in the hole with tweezers. A wooden toothpick was then stuck in the ground in the top left corner of the grid cell at 9 mm distance from the seed in the middle of the cell to facilitate localisation of the seedlings in spring. The grid was removed after sowing and each plot was then protected with a cage made out of chicken wire of 100 × 50 × 30 cm high. We thus obtained four pairs of plots, each pair with an identical composition and layout. In one of the two plots of each pair, we applied 99 g of organic fertilizer (DCM Gazonmeststof/Engrais pelouse; NPK (Mg) 9-4-7 (2)) on 24 February 2014, which gave us two replicas of four plots each, one with and one without fertilizer. The *Rhinanthus* density in each plot at sowing was 362 seeds per m2, which is relatively low compared to sowing densities used in other experiments (600-1000 m-2; Westbury & Dunnett, 2007).

Starting in March 2014, we recorded seedling emergence at least twice a week in all plots, and followed the fate of the plants until seed set. The date of emergence and the date of opening of the first flower were recorded, as well as the date of death if this happened before completion of the life cycle. For plants that survived until reproduction, we photographed the inflorescence to verify the class of the plant (*R. major, R. minor* or F_1_ hybrid) using flower morphology. We recorded the number of flowers produced (on the main inflorescence and on secondary branches if present) and harvested each fruit with the surrounding calyx using small scissors when the seeds were ripe and the capsule dehisced, storing the capsules individually in 24-well plates. The number of seeds present in each capsule was determined by removing the fruit from the well, emptying and discarding the capsule, counting the developed seeds and putting them back into the well. Some seeds may have fallen from their capsule before they could be harvested and counted (e.g. during strong winds), and some plants lost entire fruits due to herbivore damage, so the total number of seeds counted is likely to be an underestimate of the total number of seeds produced. We therefore also used the total number of flowers produced as a measure of fitness, since seed production in *Rhinanthus* is never pollen-limited (Natalis & Wesselingh, 2012a; Hargreaves *et al.*, 2015) and fruit set in the field is practically always 100% (R.A. Wesselingh, pers. obs.). A small amount of leaf material was collected for DNA extraction from each plant after flowering had finished, to minimize the impact of the removal of leaf biomass on flower and seed production. The leaf material was immediately stored at –80°C until analysis.

We checked the identity of the resulting plants for several reasons. First, *R. minor* is capable of autonomous self-pollination (Ducarme & Wesselingh, 2013) and even emasculation of a closed flower bud is not always sufficient to prevent selfing. This means that the offspring from crosses between *R. minor* and *R. major* may still contain pure *R. minor* seeds. Second, errors could have occurred during pollination, seed counting, during the transfer of the seeds from the 24-well storage plates to the 96-well plates and during sowing. Finally, because of the proximity of a mixed population of both species to the transplant site, we could not exclude that some seeds from this population would have been dispersed into the area where we had sown our experimental plots. Indeed, we did find a few plants inside the plots that were not close to a toothpick, which were considered to be intruders and excluded from our analysis. Since it is possible that other such seeds would have been present in grid cells where we had sown a seed, we checked the identity of all the flowering plants, using the photographs taken during flowering and a genetic identification tool. For this latter, we chose one species-specific SNP marker out of a panel of more than 3000 SNP markers that consistently differed between the two species we detected using ddRADseq analysis on 57 plants of the two species (K. Mirzaei & R.A. Wesselingh, in prep.) from the same source populations as the ones used to create the F_1_ hybrids in this experiment. Primers were developed to amplify the specific fragment containing the SNP using PCR, and the presence/absence of the SNP was detected by digesting the extracted and amplified DNA with an enzyme with a restriction site that contained the SNP marker. The fragment was only digested when the SNP marker for *R. major* was present, which led to an electrophoretic banding pattern with only one, undigested band of 250 bp for *R. minor*, a pattern with two bands of 70 and 180 bp, respectively, for *R. major* and a pattern with all three bands present for the F_1_ hybrids (see Supplementary information for details).

### Data analysis

All statistical calculations were done in R version 3.5.0 using RStudio version 1.1.453.

To describe the germination process under controlled conditions, we used a three-parameter log-logistic model F(*t*) = *d*/(1 + exp[*b*{log(*t*)-log(*t*50)}]), in which *t* is time (in days), *d* is the final germination percentage, *t*50 the time point at which half of the seeds have germinated, and *b* the slope of the curve at time point *t*50 (Ritz *et al.*, 2013). The model was fitted to the data for each class separately using the R package drc (Ritz *et al.*, 2015).

For greenhouse performance, we used total flower production (log10-transformed) as the dependent variable and tested for differences among classes using a linear model.

Differences among the classes in emergence and survival until flowering in the field were analysed using logistic regressions with emergence/survival as the dependent variable and class, fertilizer application and their interaction as factors. When the class effect was significant, we used pairwise G-tests (R package RVAideMemoire), with the Hochberg correction for multiple comparisons (Hochberg, 1988). Differences among the classes in the date of emergence and flowering were analysed using linear models with date as the dependent variable and class, fertilizer application and their interaction as factors. Post-hoc Tukey tests were performed using the R package emmeans when the effect of one or more factors was significant. We applied the same method to the total number of flowers and the total number of seeds produced per plant; these variables were log10-transformed first to obtain normality. In order to compare our results with those of Kwak (1980), who used the number of seeds per flower, we also analysed the per-plant average number of seeds per flower in a linear model with class and total number of flowers as factors.

## Results

### Germination under controlled conditions

The hybrid seeds formed on *R. minor* (F_1m_) were the first to start germinating and this class also reached the highest germination rate (Table 1, Fig. 1). It took only 49 days for this hybrid class to reach half of its final germination percentage, compared to 61 days for *R. minor* and 74 for both *R. major* and the F_1a_ hybrid. Only 15% of the F_1a_ hybrid seeds germinated, compared to 80% and higher for the seeds of the other three classes.

**Table 1.** Parameter estimates (standard errors in parentheses) for the germination curves of the four classes of seeds under controlled conditions.

| Class | *b* | *d* | *t*50 |
| --- | --- | --- | --- |
| *R. minor* | −14.431 (1.918) | 0.805 (0.045) | 61.17 (0.957) |
| F<sub>1m</sub> | −8.520 (0.749) | 0.981 (0.013) | 49.38 (0.998) |
| F<sub>1a</sub> | −10.185 (2.187) | 0.151 (0.035) | 74.13 (3.267) |
| *R. major* | −14.812 (1.604) | 0.861 (0.041) | 74.84 (1.159) |

**Fig. 1.**
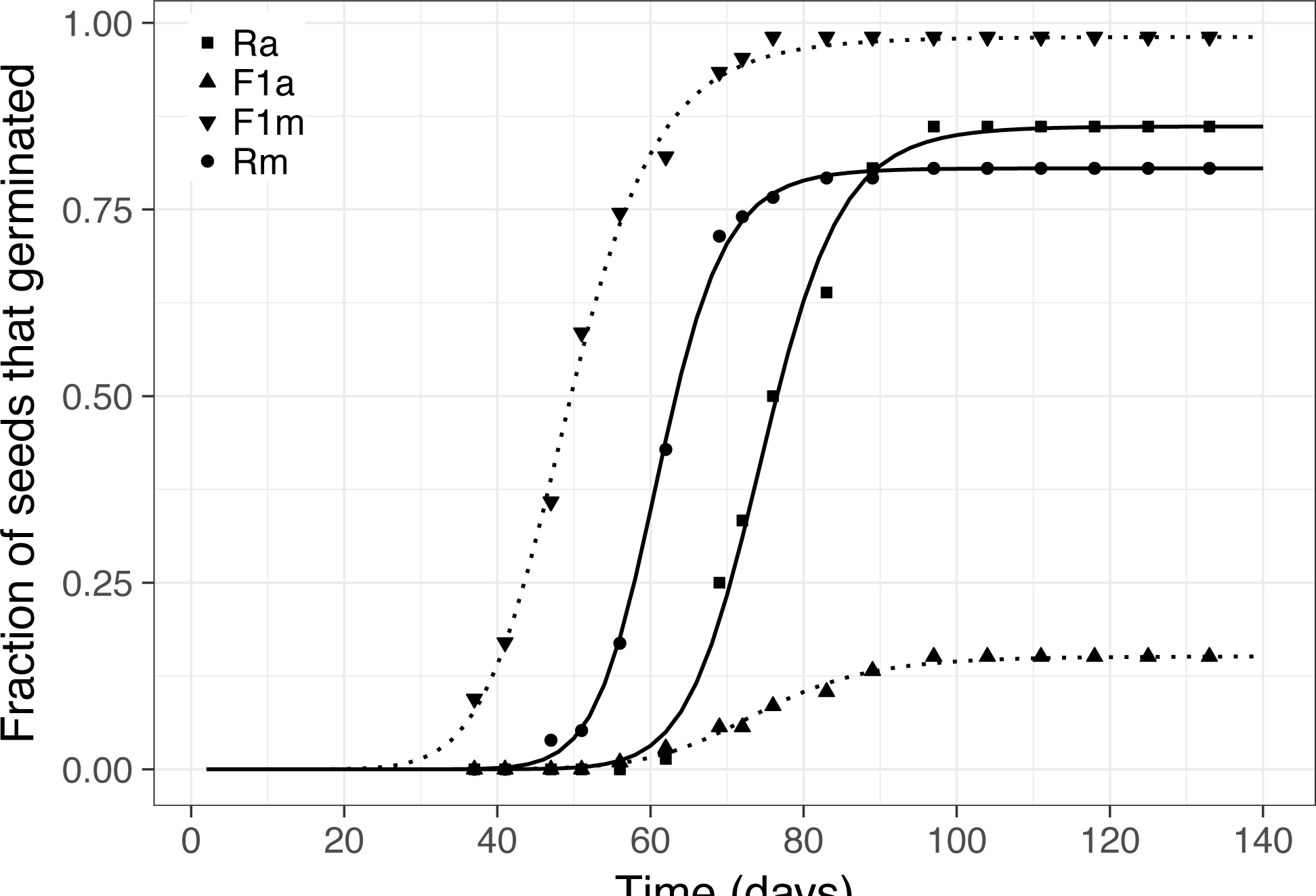
Germination over time under controlled conditions in the laboratory (5°C) with the fitted three-parameter log-logistic curves for the two species (continuous lines) and their hybrids (dotted lines). Rm = *Rhinanthus minor*, F1m = F_1m_ hybrids, F1a = F_1a_ hybrids, and Ra = *Rhinanthus major*.

### Greenhouse flower production

In 2013, there were no significant differences in flower production among the classes (Table 2, Fig. 2a). In 2014, in the new greenhouse, the number of flowers per plant was lower overall and highest in the F_1m_ hybrids, followed by *R. minor* and *R. major* with the lowest flower production (Table 2, Fig. 2b). The F_1a_ hybrids showed an intermediate flower production and did not differ significantly from the other classes.

**Table 2.** ANOVA table for the linear model on the log-transformed number of flowers produced by *Rhinanthus* plants in the greenhouse in 2013 and 2014 with class as main factor.

| Factor | df | SS | MS | *F* | P (> *F*) |
| --- | --- | --- | --- | --- | --- |
| *Greenhouse 2013* |  |  |  |  |  |
| Class | 3 | 0.2804 | 0.0934 | 1.3424 | .2697 |
| Residuals | 57 | 3.9682 | 0.0696 |  |  |
| *Greenhouse 2014* |  |  |  |  |  |
| Class | 3 | 3.3642 | 1.2081 | 14.4080 | <.0001*** |
| Residuals | 222 | 18.6139 | 0.0839 |  |  |

**Fig. 2.**
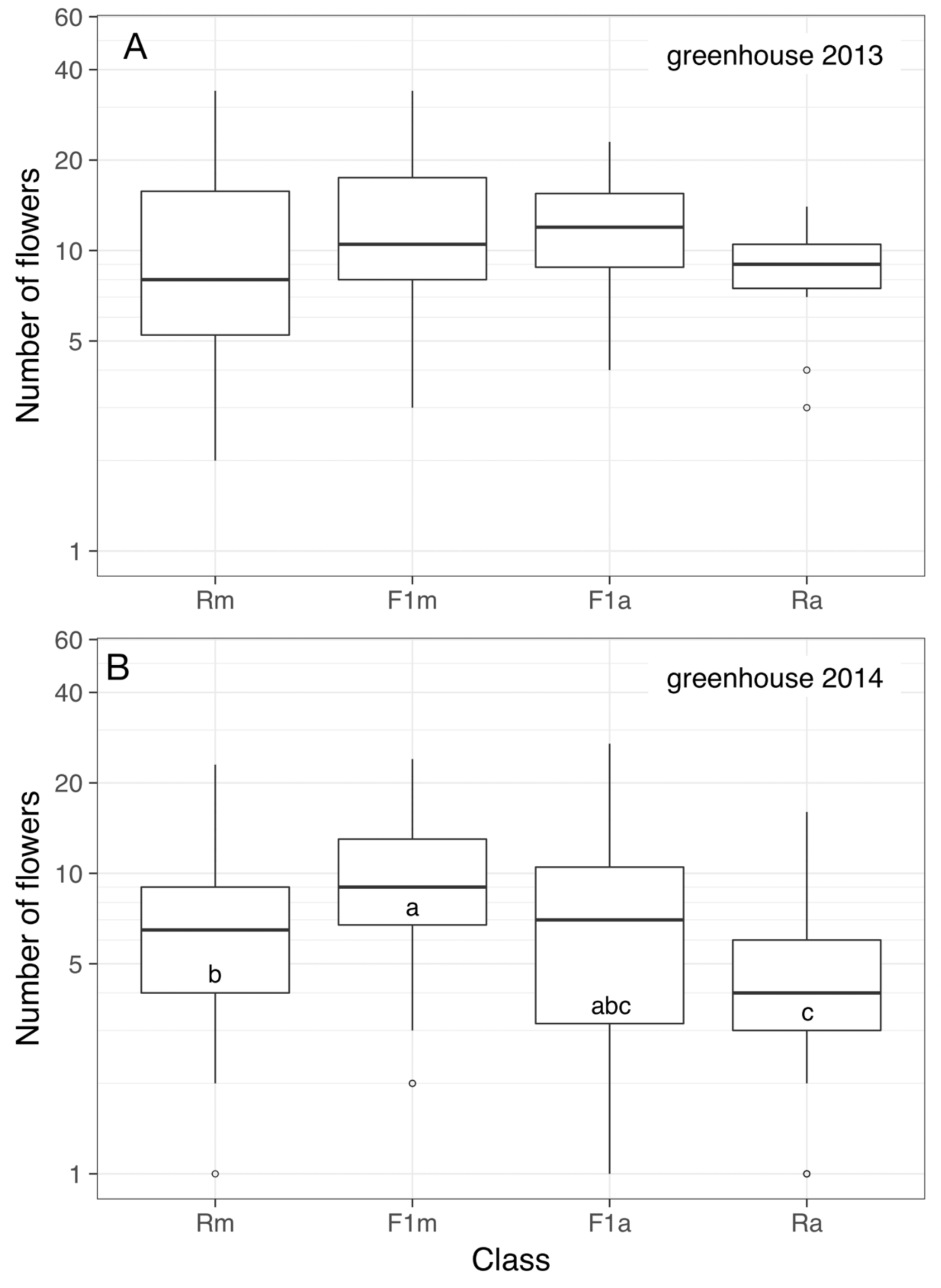
Box and whisker plots for the total number of flowers produced (on a logarithmic scale) for four classes (Rm: *Rhinanthus minor*, F1m: F_1m_ hybrids, F1a: F_1a_ hybrids, Ra: *Rhinanthus major*) in the old greenhouse in 2013 (a) and the new greenhouse in 2014 (b). Boxes that share an identical letter within each fertilizer treatment did not differ significantly from each other in post-hoc Tukey tests.

### Emergence and survival in the field

The first emerging seedlings were observed in the outdoor plots on 10 March 2014, and a total of 260 seedlings emerged at the grid positions. Of these seedlings, a total of 133 survived until flowering. Four plants were subsequently identified as intruders and excluded from the data set: two sown seeds were supposed to be F_1a_ hybrids, but the resulting plants were identified as *R. minor*, both morphologically and genetically. One *R. major* plant appeared where an *R. minor* seed had been sown, and one F_1_ hybrid emerged and flowered at the location of an *R. major* seed. Six seeds from *R. minor* x *R. major* crosses, which were expected to be F_1m_ hybrids, turned out to be (selfed) *R. minor* seeds, and these were kept in the data set and classified as belonging to the *R. minor* class. Similarly, two cases were discovered in which an F_1a_ hybrid turned out to be *R. major*, and we classified these two plants as *R. major*. Three plants died shortly after they started flowering, so no fitness data could be recorded, which resulted in 126 flowering plants for which we had at least the total number of flowers produced.

The overall emergence rate was 22.3%, and we observed some differences among the classes in emergence rate (Fig. 3), especially in the plots with fertilizer, but these were not statistically significant (Table 3). Likewise, the emergence rate was usually higher in the unfertilized plots compared to the fertilized plots, but this effect did not reach statistical significance either, nor did the interaction between class and fertilizer application, although *R. major* showed a tendency towards a higher probability of emergence in fertilized plots, in contrast to the other three classes.

**Table 3.**
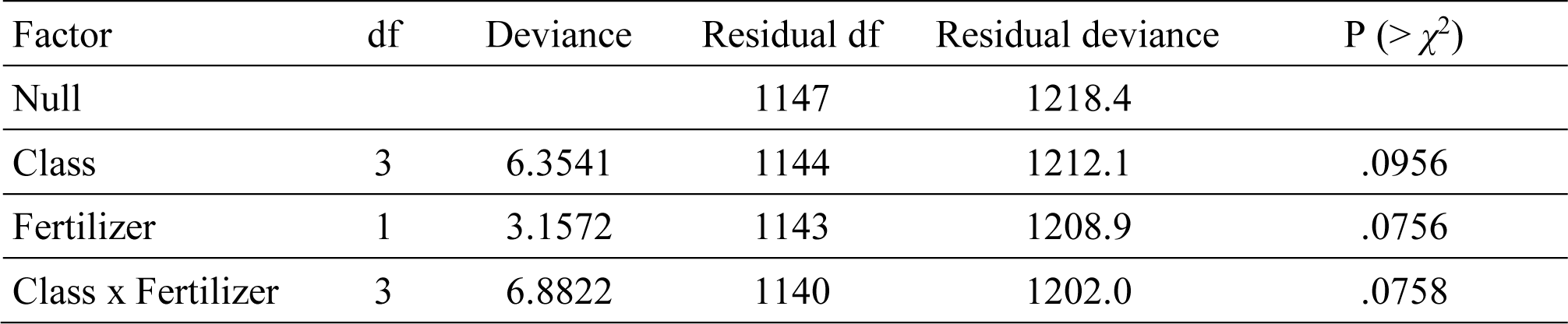
Analysis of deviance for the logistic model on the probability of *Rhinanthus* seedling emergence in the experimental field plots with class and fertilizer application as main factors.

**Fig. 3.**
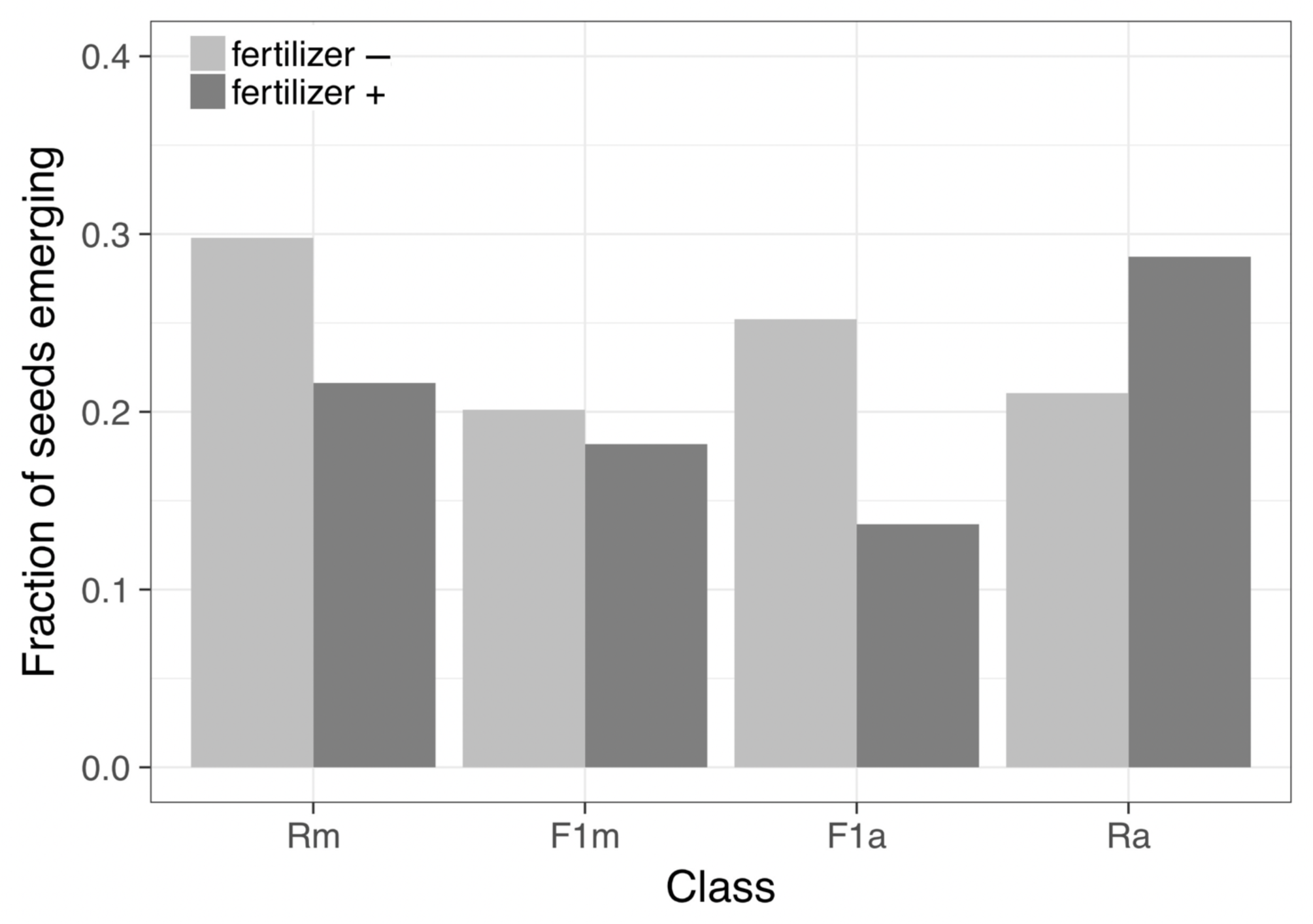
The fraction of seeds emerging as seedlings in the outdoor plots for four classes (Rm: *Rhinanthus minor*, F1m: F_1m_ hybrids, F1a: F_1a_ hybrids, Ra: *Rhinanthus major*) and two fertilizer treatments.

The date of germination differed significantly among classes as did the response in the different classes to fertilizer treatment (Fig. 4, Table 4). The F_1m_ hybrids emerged earlier than most other classes, while the F_1a_ hybrids showed a later emergence in the fertilized plots.

**Table 4.** ANOVA table for the linear model on the date of *Rhinanthus* seedling emergence in the experimental field plots with class and fertilizer application as main factors.

| Factor | df | SS | MS | *F* | P ($> F$) |
| --- | --- | --- | --- | --- | --- |
| Class | 3 | 628.2 | 209.395 | 5.4955 | .0011** |
| Fertilizer | 1 | 8.1 | 8.072 | 0.2118 | .6457 |
| Class x Fertilizer | 3 | 416.5 | 138.819 | 3.6432 | .0134* |
| Residuals | 241 | 9182.9 | 38.103 |  |  |

**Fig. 4.**
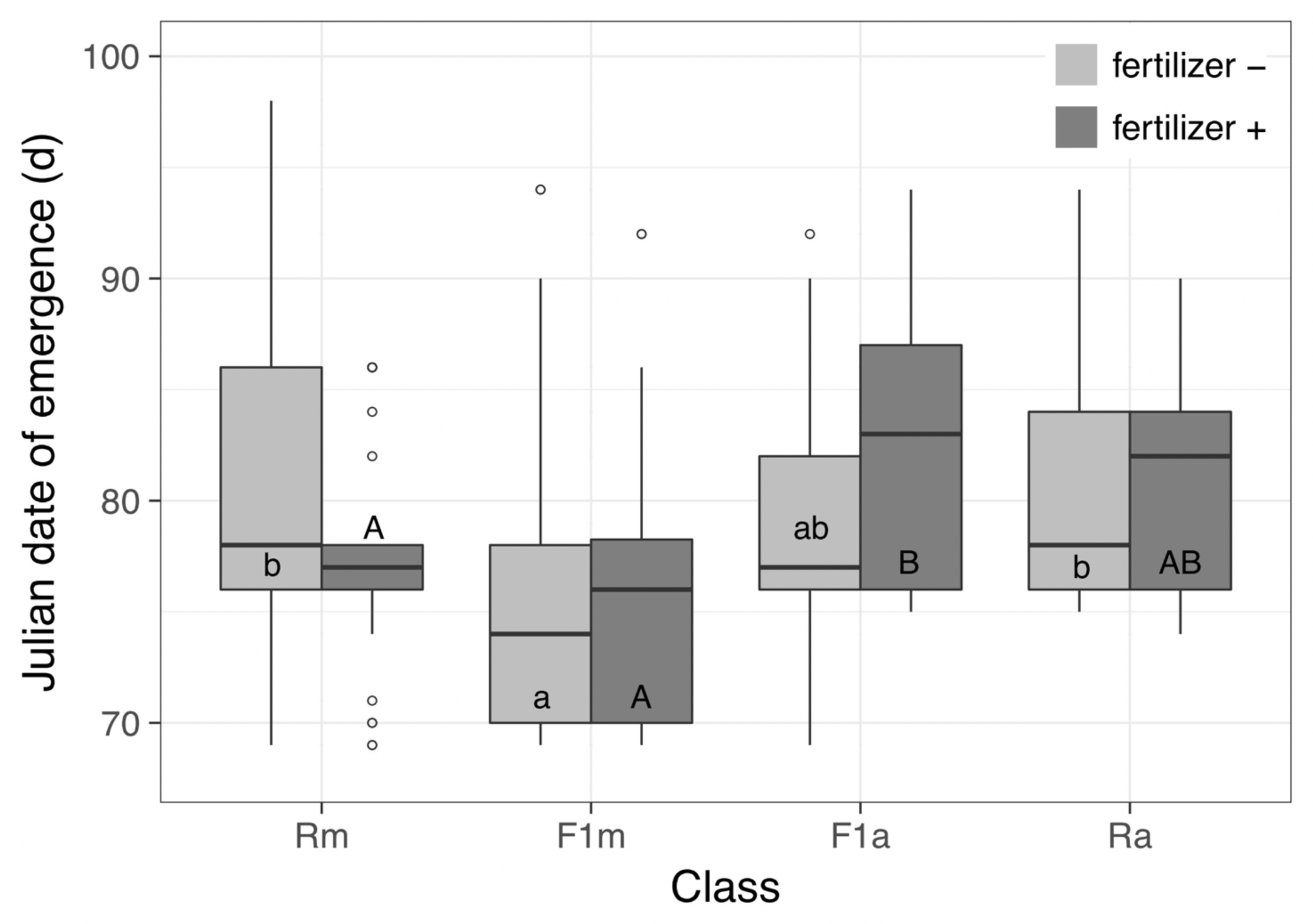
Box and whisker plots for the date of emergence, given as the Julian date, for four classes (Rm: *Rhinanthus minor*, F1m: F_1m_ hybrids, F1a: F_1a_ hybrids, Ra: *Rhinanthus major*) and two fertilizer treatments. Boxes that share an identical letter did not differ significantly from each other in post-hoc Tukey tests.

Half (50.4%) of all the seedlings survived until flowering, and there were strong differences among the classes (Fig. 5). In the unfertilized plots, *R. major* seedlings had a significantly lower survival rate than the other three classes (Table 5). Again, *R. major* survival was higher in fertilized plots compared to unfertilized plots, while this was usually reverse in the other classes, but the interaction effect was not significant. The same patterns were found when emergence and survival were combined into a single value for survival from seed until flowering (data not shown).

**Table 5.** Analysis of deviance for the logistic model on the probability of surviving from seedling to flowering with class and fertilizer application as main factors.

| Factor | df | Deviance | Residual df | Residual deviance | P ($> \chi^2$) |
| --- | --- | --- | --- | --- | --- |
| Null |  |  | 255 | 354.88 |  |
| Class | 3 | 14.0068 | 252 | 340.87 | .0029** |
| Fertilizer | 1 | 0.1017 | 251 | 340.77 | .7498 |
| Class x Fertilizer | 3 | 2.1977 | 248 | 338.57 | .5324 |

**Fig. 5.**
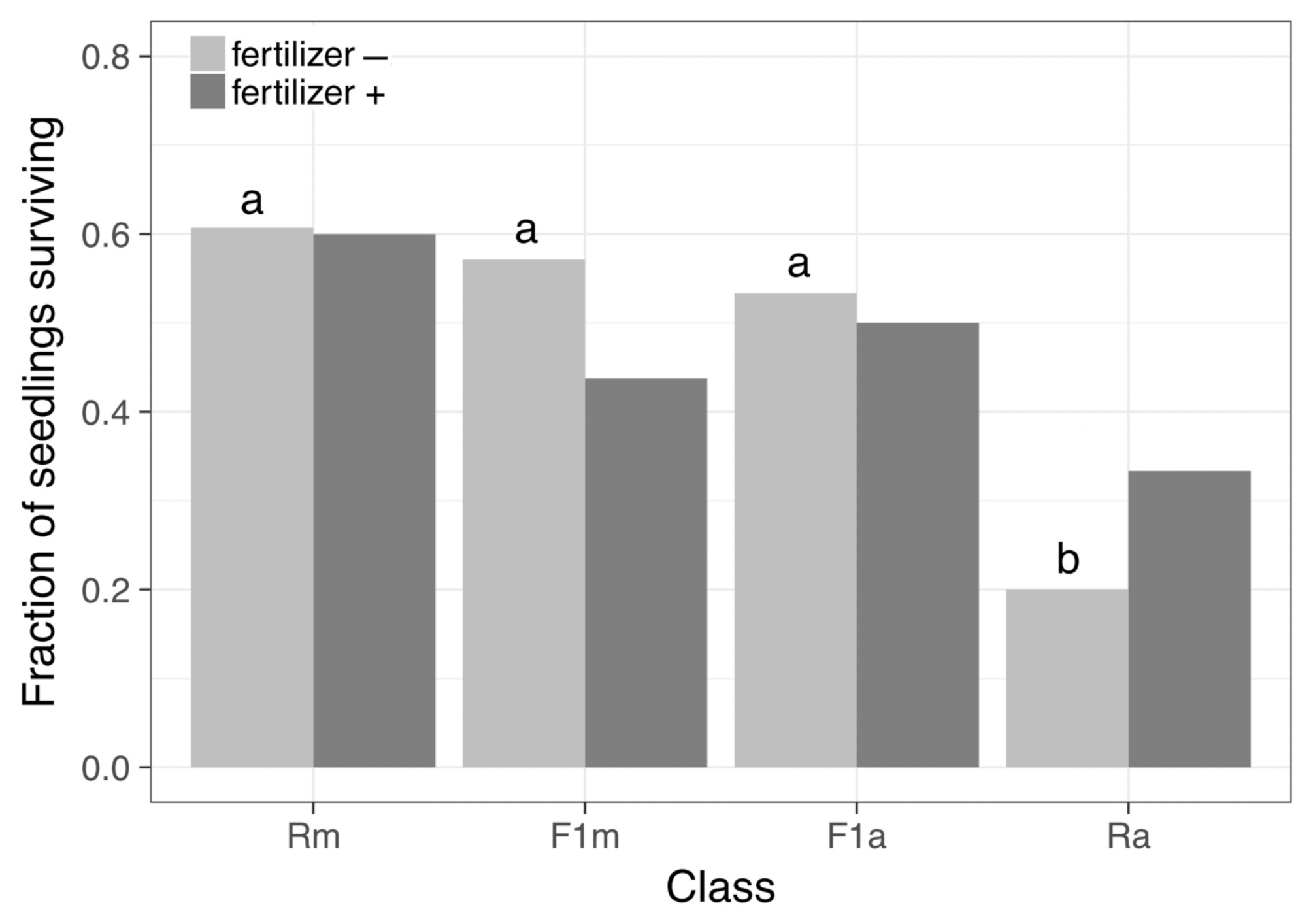
The fraction of seedlings surviving until flowering in the outdoor plots for four classes (Rm: *Rhinanthus minor*, F1m: F_1m_ hybrids, F1a: F_1a_ hybrids, Ra: *Rhinanthus major*) and two fertilizer treatments. Bars with identical letters (for the plots without fertilizer) did not differ significantly from each other in pairwise *G*-tests.

### Flower and seed production in the field

The first flower opened on 20 May 2014 and the onset of flowering was spread over six weeks. By the beginning of July, all plants but one had started flowering (Fig. 6). The F_1m_ hybrids were significantly earlier than the *R. minor* plants, and there was no effect of fertilizer application on the onset of flowering (Table 6).

**Table 6.** ANOVA table for the linear model on the date of onset of flowering of *Rhinanthus* plants in the experimental field plots with class and fertilizer application as main factors.

| Factor | df | SS | MS | *F* | P (*> F*) |
| --- | --- | --- | --- | --- | --- |
| Class | 3 | 1516.0 | 505.32 | 3.4105 | .0199* |
| Fertilizer | 1 | 115.6 | 115.64 | 0.7805 | .3788 |
| Class x Fertilizer | 3 | 144.0 | 48.00 | 0.3240 | .8080 |
| Residuals | 117 | 17335.3 | 148.16 |  |  |

**Fig. 6.**
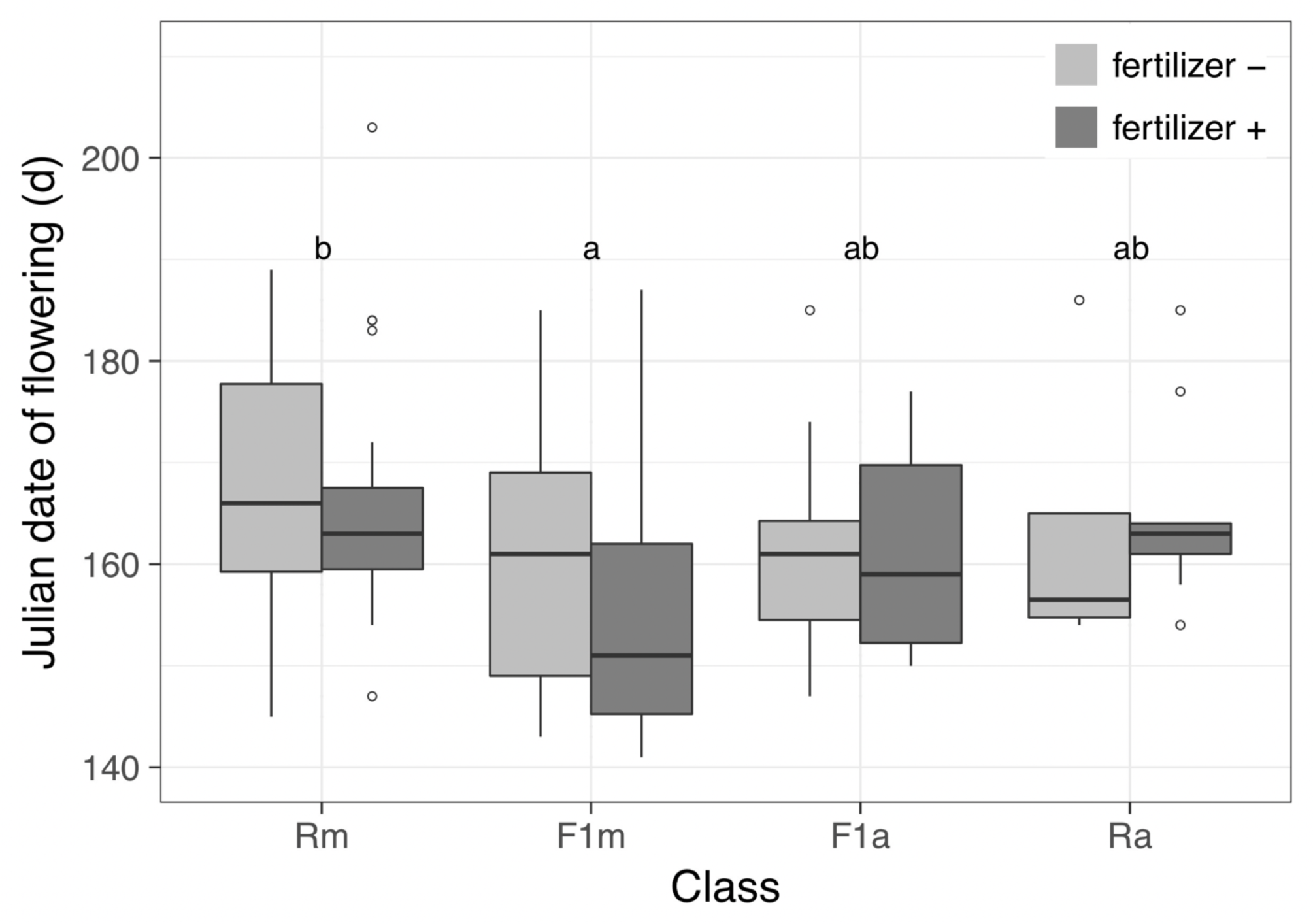
Box and whisker plots for the date of flowering, given as the Julian date, for four classes (Rm: *Rhinanthus minor*, F1m: F_1m_ hybrids, F1a: F_1a_ hybrids, Ra: *Rhinanthus major*) and two fertilizer treatments. Boxes that share an identical letter did not differ significantly from each other in post-hoc Tukey tests.

The total number of flowers produced per plant varied between 1 and 50 (Fig. 7a). There were clear differences among the classes, and flower production was much higher in the fertilized plots (Table 7). The F_1m_ hybrid class produced significantly more flowers than *R. major* without fertilizer and more flowers than both parental species with fertilizer application. The lowest number of flowers was produced by plants in the *R. major* class regardless of fertilizer treatment.

**Table 7.** ANOVA table for the linear model on the log-transformed number of flowers produced by *Rhinanthus* plants in the experimental field plots with class and fertilizer application as main factors.

| Factor | df | SS | MS | *F* | P (*> F*) |
| --- | --- | --- | --- | --- | --- |
| Class | 3 | 1.9678 | 0.6559 | 6.3167 | .0005*** |
| Fertilizer | 1 | 4.7599 | 4.7599 | 45.8389 | < .0001*** |
| Class x Fertilizer | 3 | 0.4738 | 0.1579 | 1.5210 | .2127 |
| Residuals | 118 | 12.2530 | 0.1038 |  |  |

**Fig. 7.**
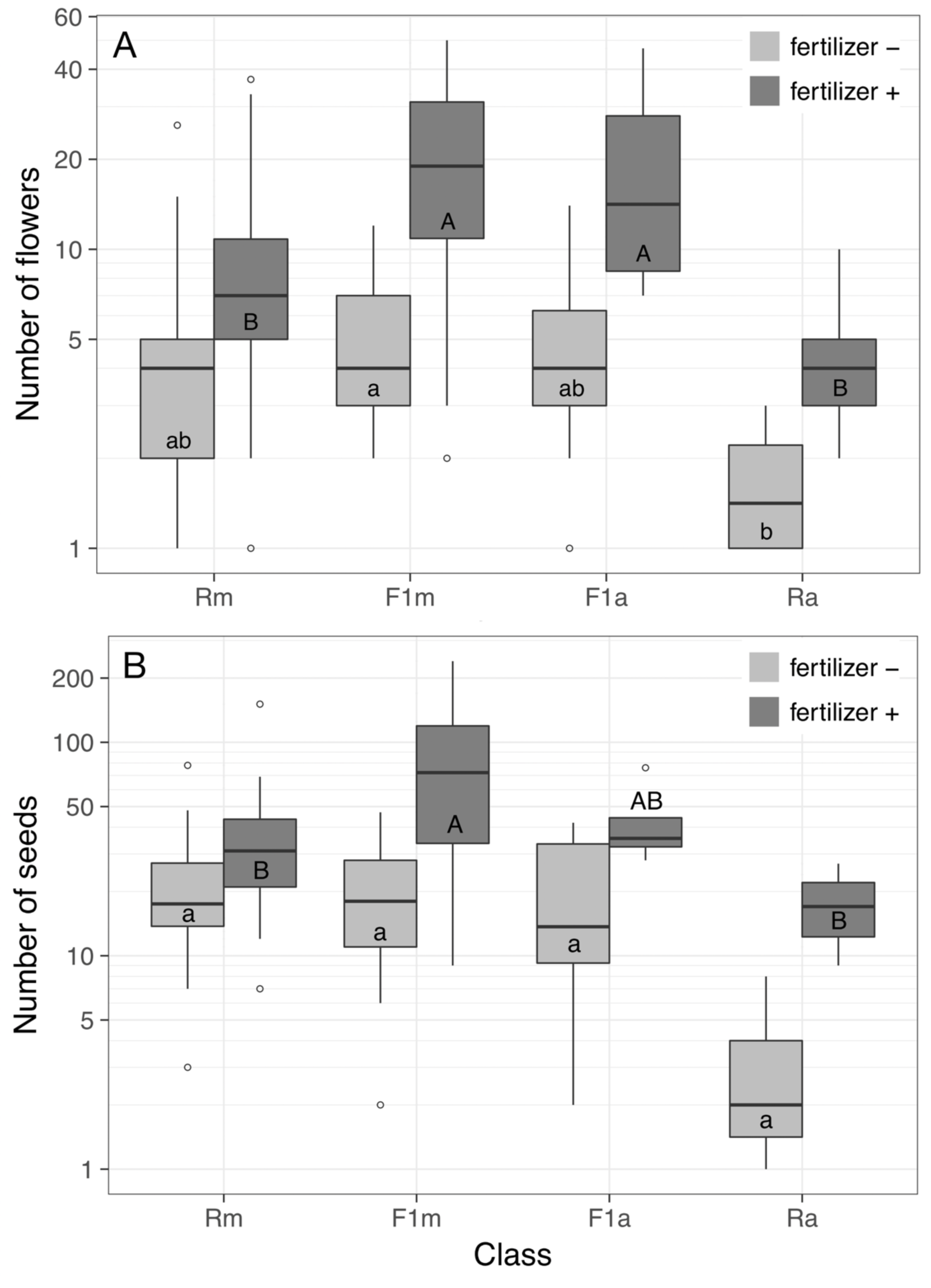
Box and whisker plots for the total number of flowers produced (on a logarithmic scale) for four classes (Rm: *Rhinanthus minor*, F1m: F_1m_ hybrids, F1a: F_1a_ hybrids, Ra: *Rhinanthus major*) and two fertilizer treatments. Boxes that share an identical letter within each fertilizer treatment did not differ significantly from each other in post-hoc Tukey tests.

The total number of seeds per plant was mainly determined by the total number of flowers per plant, which varied much more strongly among plants than the average number of seeds per fruit. Both variables together explained over 90% of the variance in seed production (linear model: seeds = –18.9086 + 4.1562 * flowers + 4.9156 * seeds per fruit, df = 96, adjusted R^2^ = 0.9173, F_2, 96_ = 544.8, P < 0.0001), but the number of flowers on its own already explained 86.65% of the variance, which increased only with an additional 5% by adding the average number of seeds per fruit to the model. A linear model with only the number of seeds per fruit explained 13.13% of the variance and adding the number of flowers to this model contributed a further 77.9% to explaining the total variance. The patterns in seed production were therefore comparable to those found when considering flower production only: an overall higher seed production in the fertilized plots and a higher seed production for F_1m_ hybrids compared to the parental classes in the plots with fertilizer (Fig. 7b).

Although quite variable among plants, the average number of seeds per flower varied much less among classes, and the linear model found no significant effect of plant class or total number of flowers (Table 8). There was a tendency for the number of seeds per flower to increase with the total number of flowers, and the nearly significant class effect was due to *R. major*, which had a steeper increase than the other classes (Supplementary figure 2).

**Table 8.** ANOVA table for the linear model on the average number of seeds per flower produced by *Rhinanthus* plants in the experimental field plots with number of flowers and class as main factors.

| Factor | df | SS | MS | *F* | P (*> F*) |
| --- | --- | --- | --- | --- | --- |
| N flowers | 1 | 7.2970 | 7.2972 | 2.8014 | .0976 |
| Class | 3 | 20.2200 | 6.7401 | 2.5876 | .0578 |
| N flowers x Class | 3 | 13.2890 | 4.4296 | 1.7006 | .1725 |
| Residuals | 91 | 237.0370 | 2.6048 |  |  |

## Discussion

### F_1_ hybrid performance

As expected from previous studies, the observed germination percentage in F_1a_ hybrids in the laboratory was much lower than for the other three classes, while the F_1m_ hybrids germinated both faster and slightly better than the parental lines. After germination, both reciprocal F_1_ hybrids between *Rhinanthus major* and *R. minor* did now show any sign of hybrid inferiority in the greenhouse: they produced as many flowers as *R. minor*, the most productive parent, and the F_1m_ hybrids actually outperformed the other parental species, *R. major*, in one year. This pattern was confirmed in the field experiment: the hybrids survived just as well as *R. minor* and outperformed *R. major* in both survival and flower/seed production in the plots without fertilizer addition. In the fertilized plots, survival was not different among the classes, but the F_1_ hybrids again outperformed *R. major* in flower/seed production, and the F_1m_ hybrids even surpassed their maternal parent. This could be a sign of heterosis (Rieseberg *et al.*, 1999), possibly caused by the fact that *R. minor* is highly selfing (Ducarme & Wesselingh, 2013), and F_1_ hybrids are more heterozygous than their maternal parent, but why this would express itself especially on the *R. minor* cytoplasmic background is not clear. In a previous study, a lower number of seeds per flower was found for hybrids (identified by flower morphology) between the two *Rhinanthus* species (Kwak, 1980). In our study, the average number of seeds per flower does not go above 6 in the F_1m_ hybrids, as it does for some of the plants in the parental lines (Supplementary figure 2), but this lower average is more than compensated for by a higher number of flowers in this class. This finding stresses the importance of measuring fitness as a whole, i.e. the total number of offspring produced, and not just a single fitness component (Arnold & Hodges, 1995).

Overall, our finding of a relatively high fitness for F_1_ hybrids is congruent with the fact that in all populations where the two parental species occur together, hybrids are found, from around 5% F_1_ hybrids in the first year (Ducarme *et al.*, 2010; Ducarme & Wesselingh, 2013) to extensive hybrid swarms, most of them close to *R. major*, in populations with a longer history of mixing (Ducarme *et al.*, 2010).

### Differences between laboratory/greenhouse and field

Our second goal was to compare performance, and especially germination, between laboratory conditions and the field situation. It turned out that the strikingly lower germination rate that has always been observed in F_1a_ hybrids in the laboratory practically disappeared under field conditions, although emergence in the plots with fertilizer tended to be somewhat lower for F_1a_ hybrids than in the other classes. We examined the data for each cross separately, and found that out of the 17 *Ra* × *Rm* crosses that were represented in the lab and in the field (a single cross was only studied in the lab and did not show any germination), all but one had a non-zero emergence rate in the field, while nine of these showed no germination in the lab. The emergence rates in the field for the crosses without germination in the lab were in the same range as those for the crosses with germination in the lab (*n* = 8). This teaches us an important lesson, which is not to rely on laboratory data only to assess hybrid fitness in our study system. Apparently, the laboratory conditions for germination, with a constant temperature of 5°C, do not sufficiently mimic outdoor conditions, where temperatures fluctuate more and seeds remain in the soil for much longer periods. Strong differences in germination rate between garden (in pots in a cold frame) and laboratory (petri dishes in a refrigerator) conditions were found by Campion-Bourget (1983) for seeds collected in pure populations of several *Rhinanthus* species.

The relative differences in timing of germination in the lab, however, are also found in the field experiment, with F_1m_ hybrids always emerging earlier than the other classes. This difference is carried over to the date of flowering, with F_1m_ hybrids again being the first to reach the flowering stage. The cold requirement for F_1m_ hybrids appears to be lower in terms of the number of cold days needed before germination, and this gives them an advantage over *R. minor*. In the greenhouse, *R. minor* develops slower than *R. major*, and an almost 3-week difference in flowering date is found between seedlings of both species planted on the same day (R.A. Wesselingh, pers. obs.). This could in part be due to a slightly lower average seed weight for *R. minor* compared to *R. major*, which will lead to slightly smaller seedlings, but there is large variation among populations and among seeds within fruits (Ernst *et al.*, 1987). It appears that *R. minor* has a lower intrinsic growth rate than *R. major*, but this has not yet been investigated systematically. In *Zea mays*, flowering time in hybrids from crosses between inbred lines was also accelerated, coupled with an increase in biomass and fertility (Birchler *et al.*, 2010).

### Effects of fertilizer addition

The addition of a single dose of organic fertilizer to half of the experimental plots in February had visible effects on the grassy vegetation: the grass became darker green and the average sward height increased from an estimated 20 cm to around 30 cm (R.A. Wesselingh, pers. obs.). The relatively nutrient-poor conditions in the unfertilized plots clearly favoured *R. minor* and both hybrid classes in the early life stages compared to *R. major*, which had a lower survival than the other three classes. The addition of fertilizer led to an increase in survival in *R. major*, while it decreased survival in the other three classes. Although the effects of class and fertilizer addition on emergence were not significant, again *R. major* reacted with an increase in emergence in the fertilized plots, while the emergence in the other three classes decreased. Flower and seed production were much higher in all classes in the plots with fertilizer. It is known that *Rhinanthus* species in general react negatively to high grassland productivity (Mudrák *et al.*, 2013), and a decrease in survival in *R. minor* as a result of fertilization of an oligotrophic meadow has been observed (Mudrák & Lepš, 2010). A positive effect of fertilizer addition on *R. major* emergence and survival in the nutrient-poor grassland in our experiment confirms the general idea that *R. major* is better adapted to more mesotrophic grasslands compared to *R. minor*.

*Rhinanthus* species can occur a diverse range of grassland habitats on different soil types, with large variation in water and nutrient availability (Westbury, 2004). Although we obtained data for the full life cycle of the two parental species and their F_1_ hybrids, our field experiment only looked at a single habitat type in a single year. This has given us valuable insight into the fitness of these first-generation hybrids, suggesting that they can perform as well as the parent with the best performance in this given situation, but more field transplants are needed to cover the full range of habitat types and to account for variability among years (Postma & Ågren, 2018). Since the formation and establishment of F_1_ hybrids are clearly not a bottleneck, we will focus our future work on advanced hybrids, including F_2_ and backcrosses, but also hybrids in natural populations, not only to determine fitness in transplant experiments, but also to identify introgressed loci involved in local adaptation (Martin *et al.*, 2006; Suarez-Gonzalez *et al.*, 2018). In our study system, hybrids close to *R. major* are much more frequent, because the pollinating bumblebees visit the hybrids as often as the more attractive *R. major*, while *R. minor* is highly selfing and less visited. This leads to unilateral introgression from *R. minor* into *R. major* (Ducarme & Wesselingh, 2005; Ducarme *et al.*, 2010), and work is currently underway to study which parts of the *R. minor* genome introgress preferentially into the *R. major* background and if they confer a fitness advantage and thus can cause adaptive introgression.

## Acknowledgements

The authors thank Marie Eve Renard for counting the seeds produced in the greenhouse and in the field and the Dr. Siebold Bargehassus Fund for generous donations to support this research. This is publication number BRC429 of the Biodiversity Research Centre of UCLouvain.

