## Supplemental material for "Fitness of reciprocal F_1_ hybrids between *Rhinanthus minor* and *Rhinanthus major* under controlled conditions and in the field"

**Supplementary file S1**

Protocol used for genetic identification of parental lines and hybrids

Genomic DNA was extracted from 0.1 g of young leaf tissue using a modified version of the CTAB protocol (Doyle & Doyle, 1990). The quality and quantity of all DNA samples were checked by using 1% agarose gels and a Qubit fluorometer (Thermo Fisher Scientific), respectively. The final concentration of each DNA sample was adjusted to 50 ng/$\mu$.

DNA amplification was carried out in a PCR reaction with a volume of 20 $\mu$l containing 50 ng genomic DNA, 2 $\mu$l of 10 X buffer, 0.4 mM dNTPs, 1.5 mM MgCl2, 2.0 U Taq polymerase, 5 pM forward primer (5'CACCCTGATTTCTCTTTCTTCAA) and 5 pM reverse primer (5'TTAAGACCCCATAAAAAGGAGGA).

DNA amplification started with 5 min at 94°C, followed by 35 cycles of 30 s at 94°C, 30 s at 52°C, 30 s at 72°C, one cycle of 5 min at 72°C and ended with storage at 4°C. 5 $\mu$l of each PCR product was electrophoresed on an 2% agarose gel at 80 V for 1.5 h and visualized under UV light. The remaining PCR product was digested in a volume of 20 $\mu$l including 5 units of *Rsa*I enzyme and 2 $\mu$l of 10 X Cutsmart buffer. The digestion procedure was performed at 37°C for 3 h without heat killing of the enzyme afterwards. Finally, digestion products were electrophoresed on 2% agarose gels at 80 V for 1.5 h and visualized under UV light. Molecular weights were estimated using a 100-bp DNA ladder on every gel.

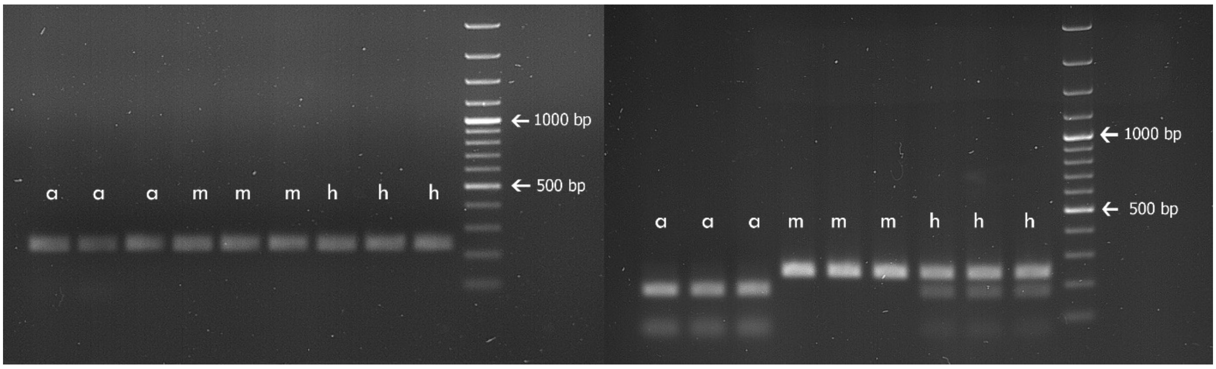


**Fig. S1.** Marker fingerprinting of three *Rhinanthus major* (a), three *R. minor* (m) and three $F_1$ hybrids (h). The picture on the left shows the 250-bp band for all the samples before digestion and the picture on the right the species-specific fragments in each sample after digestion with *Rsa*I.

**Wesselingh, R.A., Hořčicová, Š. & Mirzaei, K. Fitness of reciprocal $F_1$ hybrids between *Rhinanthus minor* and *Rhinanthus major* under controlled conditions and in the field**

**Supplementary file S2**

Number of seeds per flower in *Rhinanthus minor*, *R. major* and their $F_1$ hybrids: figures (Fig. S2) and linear model (Table S2).

**Fig. S2.** The number of seeds per flower in each of the four classes: *Rhinanthus minor* (Rm), $F_1$ hybrids with *R. minor* as the maternal parent (F1m), $F_1$ hybrids with *R. major* as thematernal parent (F1a) and *Rhinanthus major* (Ra).

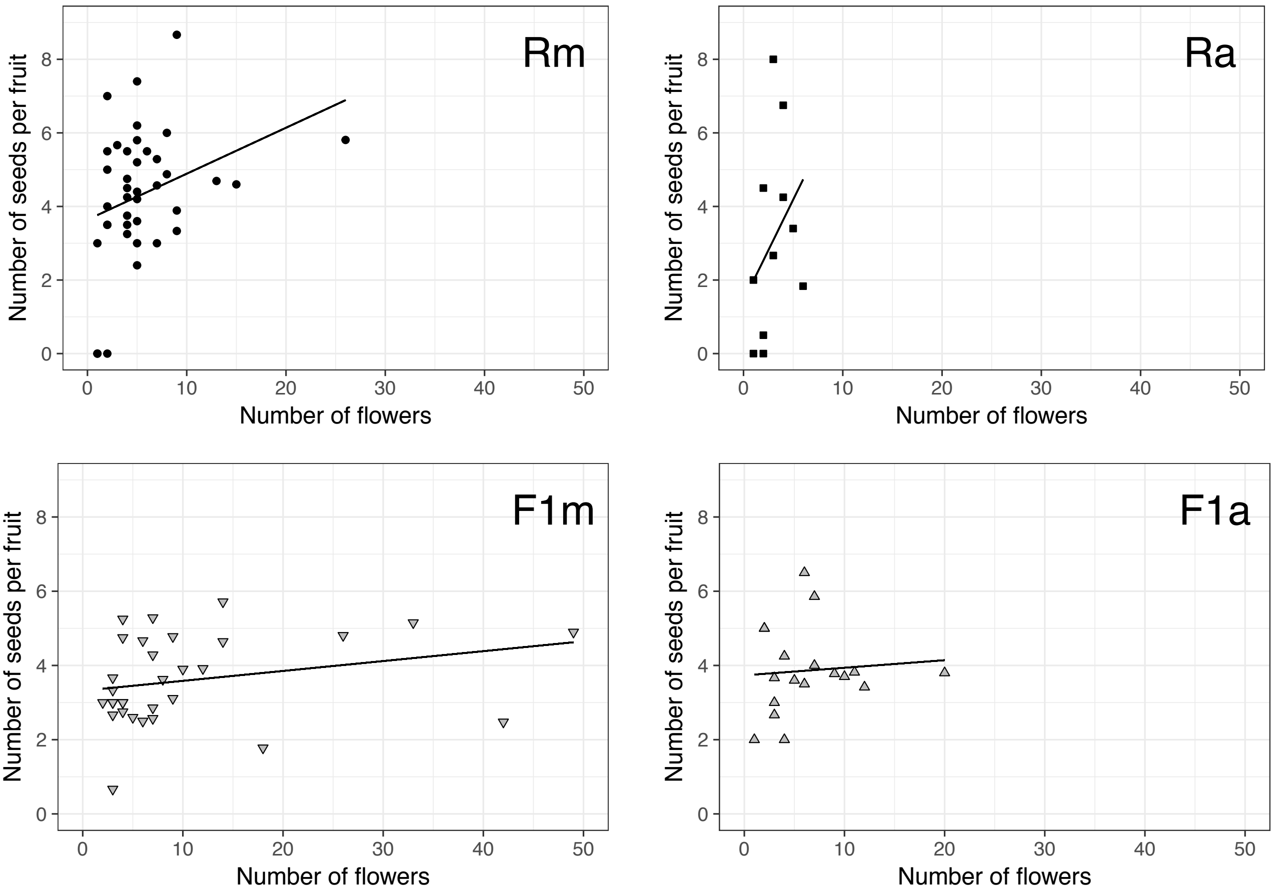


**Table S2.** ANOVA table for the linear model on the number of seeds per fruit produced by *Rhinanthus* plants with complete data (all fruits collected) in the experimental field plots with class and number of flowers as factors.

| Factor | df | SS | MS | F | P (> F) |
|---|---|---|---|---|---|
| Number of flowers | 1 | 7.297 | 7.2972 | 2.8014 | 0.0976 |
| Class | 3 | 20.220 | 6.7401 | 2.5876 | 0.0578 |
| Class x N flowers | 3 | 13.289 | 4.4296 | 1.7006 | 0.1725 |
| Residuals | 91 | 237.037 | 2.6048 | | |
